# Spatial structure affects the establishment and persistence of closed microbial ecosystems

**DOI:** 10.1101/2024.06.28.601237

**Authors:** India Mansour, Maximilian Hähnlein, Luise Minkewitz, Emely Noa Wilk, Mitja Remus-Emsermann, Janis Antonovics, Matthias C. Rillig

## Abstract

Why Earth has remained habitable for billions of years is a question that has long fostered debate in biology and earth sciences. Closed systems approaches have yielded information about the underlying mechanisms, including the persistence of matter recycling. However, the majority of these studies have been conducted under relatively homogenous conditions using aquatic systems. Here, we investigated the effect of spatial structure and heterogeneity on the persistence or failure of closed microbial biospheres. This mimics unsaturated soil-like conditions that were inoculated with a two species producer-decomposer community. Specifically, we investigated how microhabitat physical structure and necromass spatial distribution affected population dynamics and system time-to-failure. We observed strong effects of microhabitat structure, including particle size and moisture level, on persistence at both the population and system levels. Systems containing the smallest substrate particles failed quickly and on average did not support decomposer populations except at high initial cell densities. Persistence was promoted by larger substrate particles, likely due to larger pore sizes resulting in shorter movement distances and better accessibility to resource patches (i.e. necromass). Building on these findings, we manipulated necromass patch distribution and observed that algae clustered around necromass patches when present. Necromass patch distribution had small but significant effects on persistence, with lower persistence in intermediate vs. high or low necromass heterogeneity. Together these findings indicate a limit to the spatial/physical parameter space in which producer-decomposer communities can establish and self-sustain via self-recycling of necromass.

## INTRODUCTION

Earth is a largely matter-closed and energy-open biosphere that has sustained life for billions of years (Betts et al., 2018). Biological activity has changed Earth’s surface and atmosphere, and these changes in turn affected biological activity and evolution (Gutiérrez-Preciado et al., 2018). Biological-earth system feedbacks (e.g. biological modification of the atmosphere and environmental species filtering) have been posited as responsible for global biodiversification events (Smith and Harper, 2013) and for maintaining habitable conditions capable of supporting ecological assemblages for up to millions of years (DiMichele et al., 2004). The regulatory processes that sustain life on Earth are themselves driven by biological actors, a concept famously explored by Lovelock and Margulis in the Gaia hypothesis (Lovelock and Margulis, 1974). While controversial, their idea of Earth as a self-regulating superorganism has sparked biological debate, with attempts to reconcile the Gaia hypothesis with Darwinian principles and explain the persistence of matter recycling (Doolittle and Inkpen, 2018; Boyle and Lenton, 2022). These works all approach the same central question: how do life and life supporting systems persist over time?

Short of being able to replicate the planet itself, closed ecosystems approaches are well-suited to explore questions of biological persistence and matter recycling (Taub, 1974; Kearns and Folsome, 1981; Obenhuber and Folsome, 1988; Rillig and Antonovics, 2019), and have been proposed as a means to investigate global carbon cycling (Milcu et al., 2012). Modeling closed systems has furthered our understanding of self-sustaining systems, including thermodynamic restraints on matter recycling (Goyal et al., 2023). Experimental studies employing closed systems have led to insights about microbial self-recycling (Rillig et al., 2021; Shoemaker et al., 2021) and multi-species self-organization of carbon and nutrient cycling (Obenhuber and Folsome, 1988; de Jesús Astacio et al., 2021). Some studies have focused more specifically on producer-decomposer relationships in closed systems, providing evidence for coexistence without explicit conditions preventing competitive exclusion (Daufresne et al., 2008) and even the emergence of a producer-decomposer mutualism (Hom and Murray, 2014).

The majority of such studies have been conducted in aquatic environments, such as continuously homogenized complex nutrient media. However, microorganisms more commonly inhabit and/or create spatially structured environments (Vos et al., 2013; Cremin et al., 2023), which has important consequences for species interactions and resource diffusion/distribution (Dechesne et al., 2008; Kovács, 2014; Duxbury et al., 2024). Furthermore, most environmental microorganisms live in energy and nutrient-limited conditions (Lever et al., 2015). In soils, for example, resources are heterogeneously available in both space and time. We here studied microbial population and system persistence times in spatially structured and energy-limited closed systems (i.e. microbial biospheres), to investigate factors affecting rates of system persistence (Rillig and Antonovics, 2019).

We hypothesized that substrate particle type and size would affect system persistence (operationally defined as a detectable chlorophyll autofluorescence signal from closed units) through modulating organismal proximity to resources and each other. To test this hypothesis, we manipulated spatial structure and resource heterogeneity in closed, necromass-recycling-based systems and investigated microbial population and system persistence. In total, we conducted four microbial biospherics experiments that tested varying substrate type and size (1-3), liquid volume (2) and resource heterogeneity (4).

## RESULTS

### Experiment 1: Substrate size and type affects microbial biosphere persistence time

In our first experiment, we explored the effects of different substrate particle types (1-2 mm sand, 1.0 and 0.5 mm glass beads and no substrate particles) on the persistence of microbial biosphere systems over 19 days, when the chlorophyll autofluorescence signal was lost from about half of the biospheres. While biospheres containing sand or no substrate (liquid) maintained a chlorophyll autofluorescence signal during the entire incubation period (Fig. 1a), a significant difference was observed in failure time between biospheres containing 0.5 mm and 1.0 mm glass beads (log rank test *X*^2^= 18.1, df = 1, p < 0.0001). The median survival time for biospheres with 0.5 mm beads was lower than for 1.0 mm beads: 10 d (95% CI 10-Inf), compared to 14 d (95% CI 14-17), respectively (Fig. 1b). After 19 days of incubation, biospheres were destructively harvested and cell densities determined. Both *E. coli* (viable) and *C. reinhardtii* (total) population densities were the lowest in biospheres containing small beads and highest in those containing large beads (Dunn test with Bonferroni correction, p < 0.05, Figure S1).

**Figure 1.**
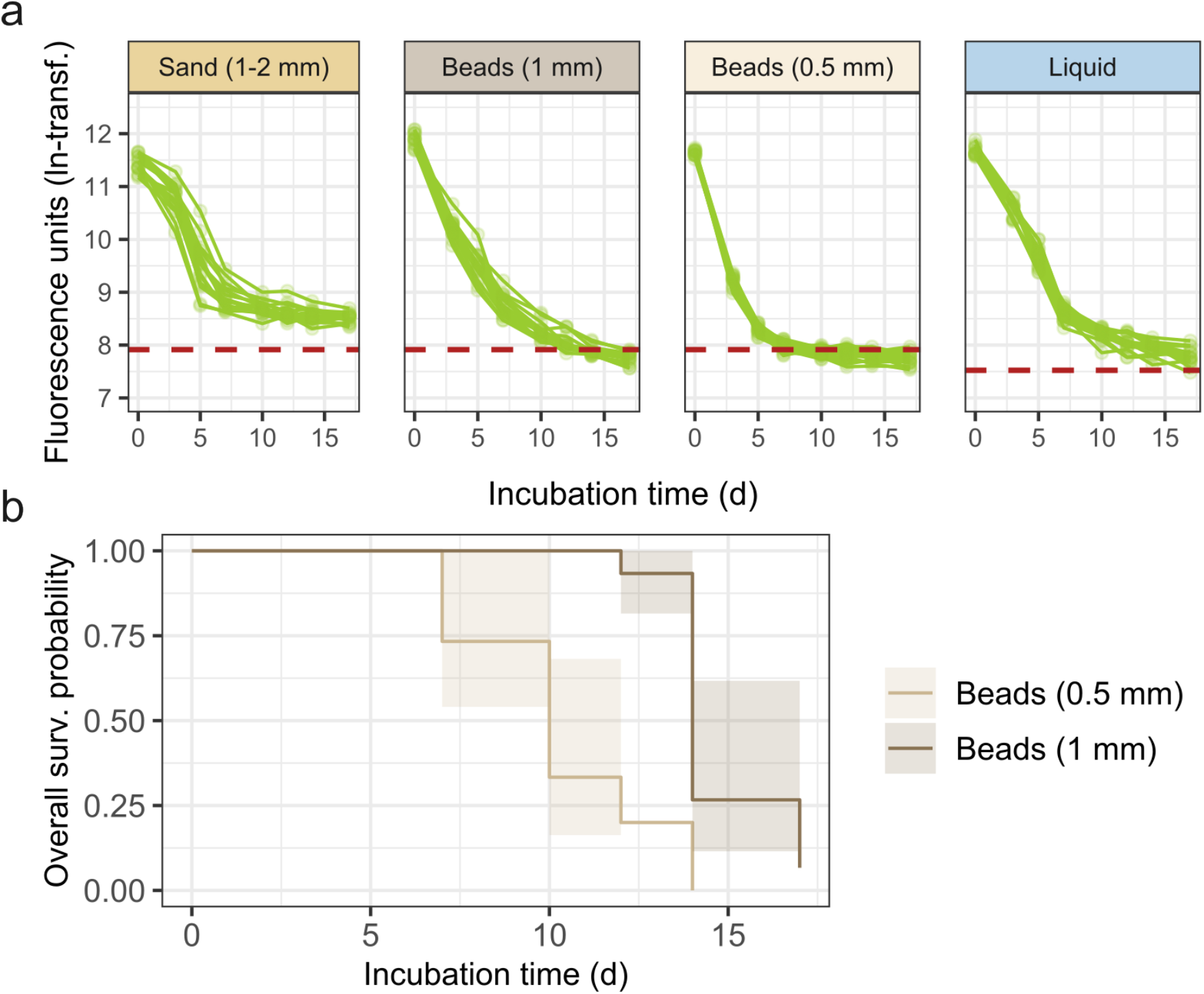
(a) Chlorophyll autofluorescence over time; each line represents the natural log-transformed autofluorescence measured for a single biosphere (n = 15 per treatment). The red dashed line indicates the threshold for system failure (i.e. the detection limit); background fluorescence was lower in biospheres with no substrate (liquid only) therefore that treatment has a lower threshold value. (b) Kaplan-Meier plot showing the overall survival probability of biospheres for treatments where a sufficient number of units failed to allow us to perform survival analysis (i.e. those containing 0.5 mm and 1.0 mm glass beads). Biosphere persistence time varies by spatial structure/substrate particle type.

### Experiment 2: Substrate particle size affects biosphere failure rate and decomposer persistence, moisture level affects biosphere persistence time

Upon observing the effects of substrate particle size on biosphere persistence, we set up a follow-up experiment investigating the effect of substrate particle size in more detail, in combination with the ‘total explorable space’ by modifying moisture level. Because large differences were observed in the glass bead treatments in experiment 1, we applied these same treatments, as well as a ‘mixed bead’ treatment, containing equal weights of 1.0 and 0.5 mm glass beads. We also applied two moisture level treatments by differing the total volume of saline in the biospheres (100 and 200 μL). We used larger well plates (48 vs. 96-well) and inoculated them with 2.5 × 10^6^ *C. reinhardtii* and 2.5 × 10^8^ *E. coli* per biosphere (microtiter well). This reduced the cell density per unit surface area in each biosphere compared to those used in experiment 1 by a factor of 2.11.

The chlorophyll autofluorescence signal from all biospheres containing small and mixed bead treatments dropped precipitously within the first 3 days of the experiment, indicating a rapid decline of algal viability (Fig. 2). Few biospheres containing 1.0 mm beads failed (n = 3/24) during the incubation period, while those containing either 0.5 mm beads or the bead mixture failed within the first two weeks (Fig. S3). Biospheres with higher moisture levels failed faster: those containing 0.5mm beads and a bead mix with 100μL failed after 14d (95% CI 12-Inf) and 13d (95% CI 12-Inf), respectively (median survival time), while those containing 200μL failed after 9.5 (95% CI 6-Inf) and 7 (95% CI 5-Inf) days, respectively.

**Figure 2.**
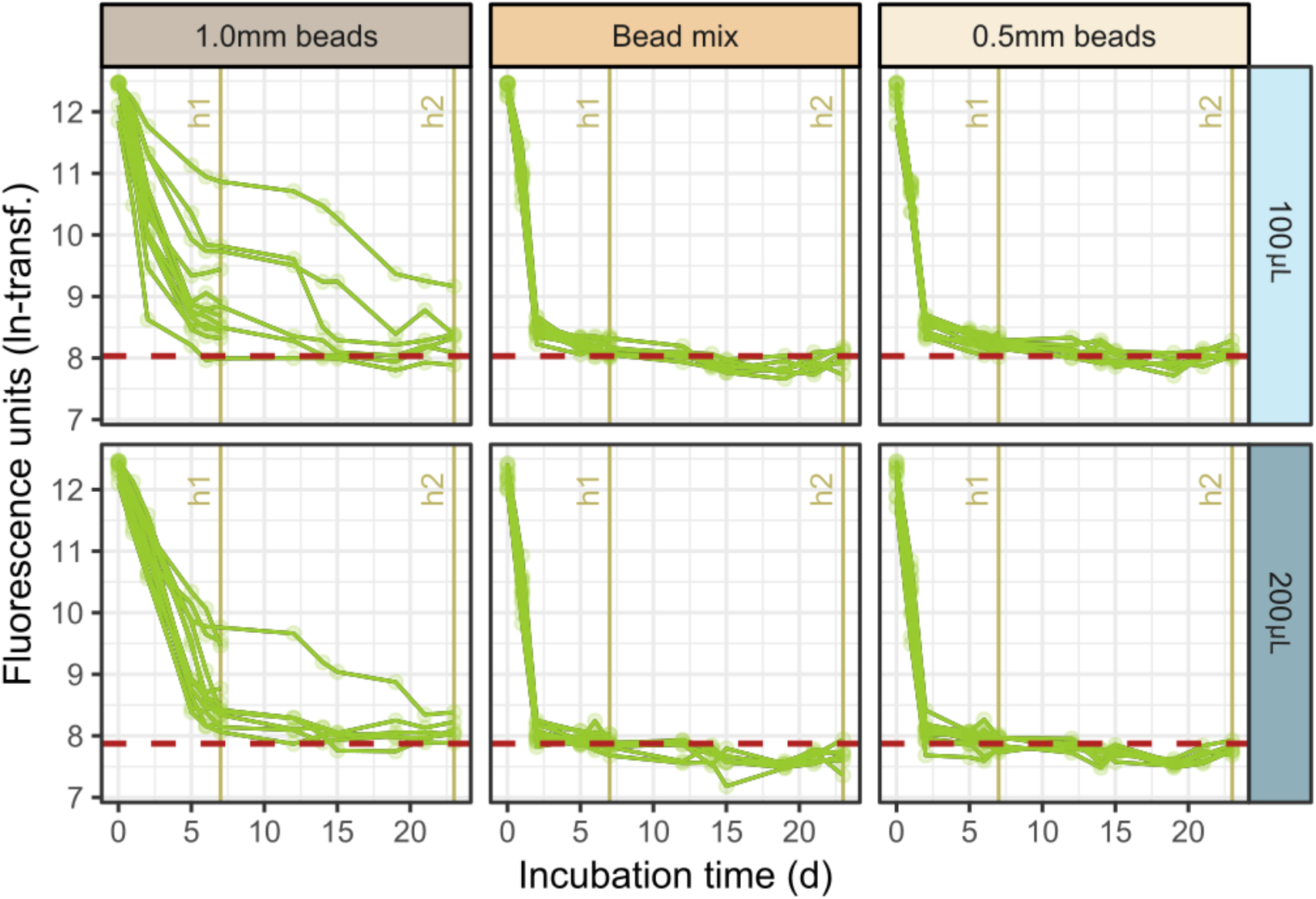
Chlorophyll autofluorescence over time; top row = biospheres containing 100μL, bottom row = biospheres containing 200μL. Each line represents the natural log-transformed autofluorescence measured for a single biosphere (n = 12 per treatment). The red dashed line indicates the threshold for system failure (i.e. the detection limit); background fluorescence was lower in biospheres with 200μL volumes, therefore that treatment has a lower threshold value. Two destructive harvests were conducted on days 8 and 29, the last measurement date before each harvest is indicated with a vertical line (h1 = harvest 1; h2 = harvest 2). Biospheres fail faster with smaller substrate size and higher moisture levels.

Microbial population densities were enumerated at two destructive harvests after 8 and 29 days of incubation. After 8 days, few viable *E. coli* were recovered from the mixed bead treatments and the 0.5 mm bead treatments (ranging below the detection limit to thousands of cells per biosphere, except one biosphere in the 0.5 mm bead treatment) but were present in all biospheres containing 1.0 mm beads (median cell density = 9.6 × 10^7^ cells/biosphere). Substrate particle size (F = 37.019, df = 2, p < 1e-07) and moisture level (F = 31.786, df = 1, p < 1e-05) both significantly affected *E. coli* population densities after 8 days of incubation. By day 29, viable *E. coli* were only recovered from the 1.0mm bead treatment and in some cases the population density even exceeded the initial inoculum (Fig. 3).

**Figure 3.**
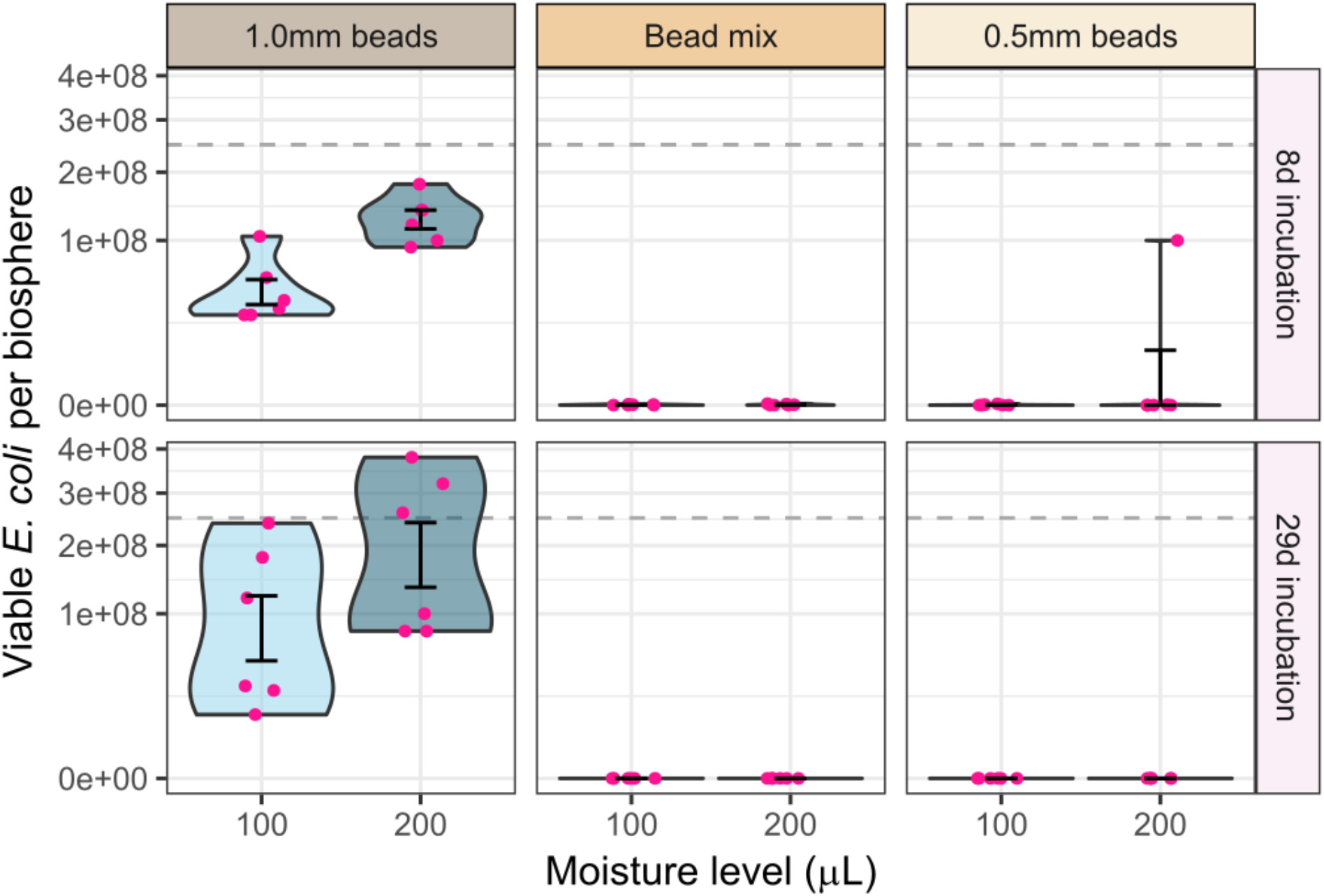
Colony forming units (CFU) were assessed via plating after 8d (top row) and 29d (bottom row) of incubation. Two plates containing three replicates of each treatment per plate (total n = 6) were destructively harvested at each time point. The gray horizontal dashed line indicates the initial cell density at the time of inoculation. Error bars show mean ± standard error. *E. coli* population densities dropped rapidly in biospheres with small-sized substrate, and were maintained in biospheres containing only larger substrate particles (1.0 mm beads).

### Experiment 3: Substrate particle type has consistent effects on biosphere failure time and rate regardless of algal species

Following up on the notable loss of viable *E. coli* populations in all 0.5 mm and mixed bead treatments observed in experiment 2, we replicated some of those treatments (1.0 and 0.5 mm glass beads with 200 μL), again using the *C. reinhardtii* and *E. coli* (CR + EC) community. We also inoculated some biospheres with a *C. moewusii* and *E. coli* (CM + EC) community to test if the observed dynamics would repeat with a different algal species. We hypothesized that the *E. coli* population decrease in biospheres containing 0.5 mm beads resulted from limited (necromass) resource access in that treatment, so as in experiment 1 we included a “liquid” treatment here, i.e. only saline and no solid substrate particles.

We observed a significantly longer persistence time for all treatments containing 1.0 mm beads compared to 0.5 mm beads: 35 d vs. 6 d for the CR + EC community (log rank test *X*^2^ = 39, df = 1, p < 10^−9^) and 21 d vs. 10 d for the CM + EC community (log rank test *X*^2^ = 39, df = 1, p < 10^−9^; Fig. 4). Furthermore, viable *E. coli* persisted in all 1.0 mm bead and liquid-only biospheres during the duration of the experiment, while viable *E. coli* densities were below the detection limit in the 0.5 mm bead biospheres by day 14 regardless of the primary producer (Fig. 5).

**Figure 4.**
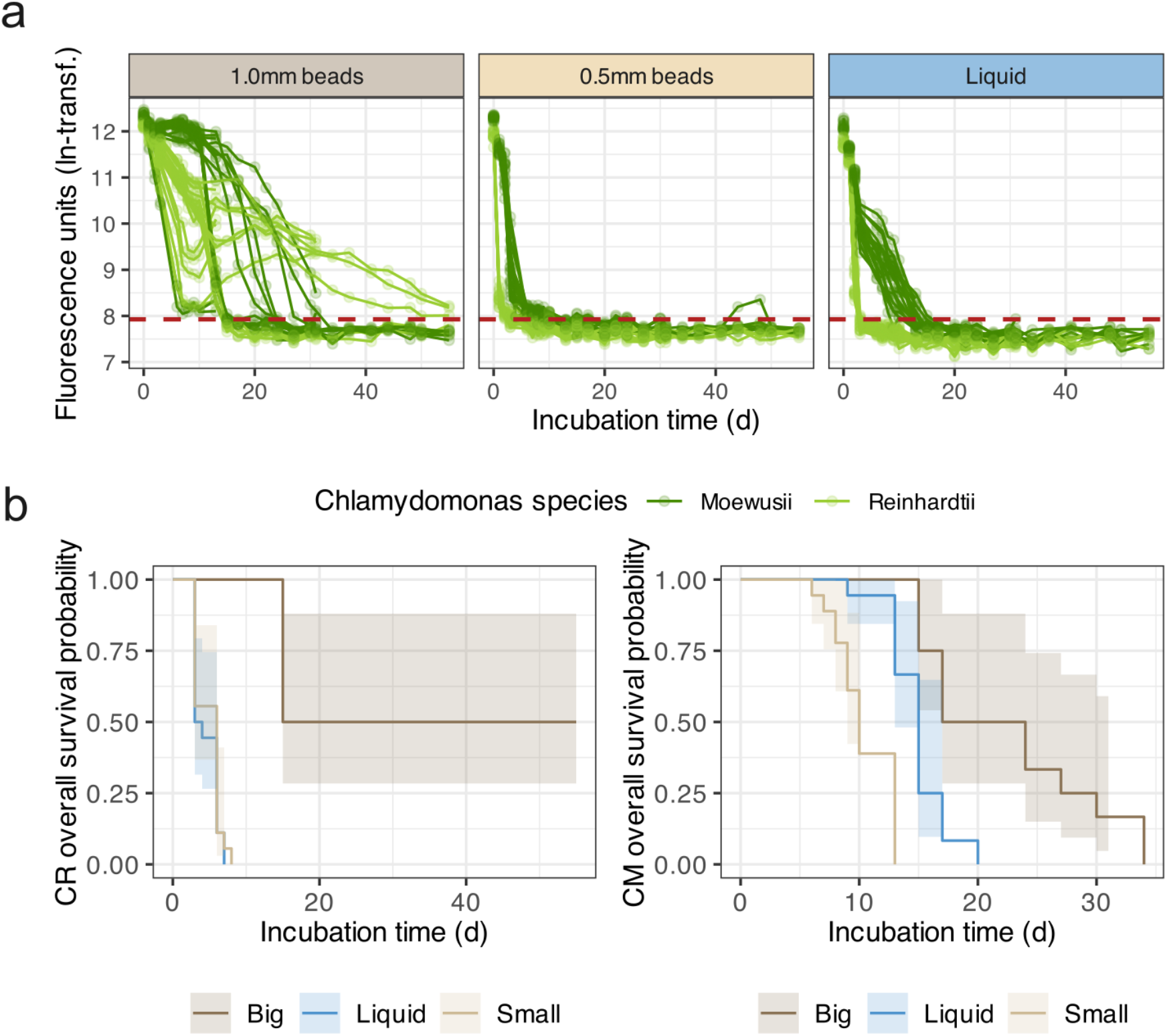
(a) Chlorophyll autofluorescence over time; each line represents the natural log-transformed autofluorescence measured for a single biosphere (n = 18 per treatment). Curves for biospheres containing *Chlamydomonas moewusii* are plotted in dark green and those with *Chlamydomonas reinhardtii* in light green. The red dashed line indicates the threshold for system failure (i.e. the detection limit). (b) Kaplan-Meier plots showing the overall survival probability of biospheres for treatments for biospheres containing *C. reinhardtii* (L) and *C. moewusii* (R) as the primary producer. Biospheres fail more quickly with smaller substrate particle size regardless of primary producer.

**Figure 5.**
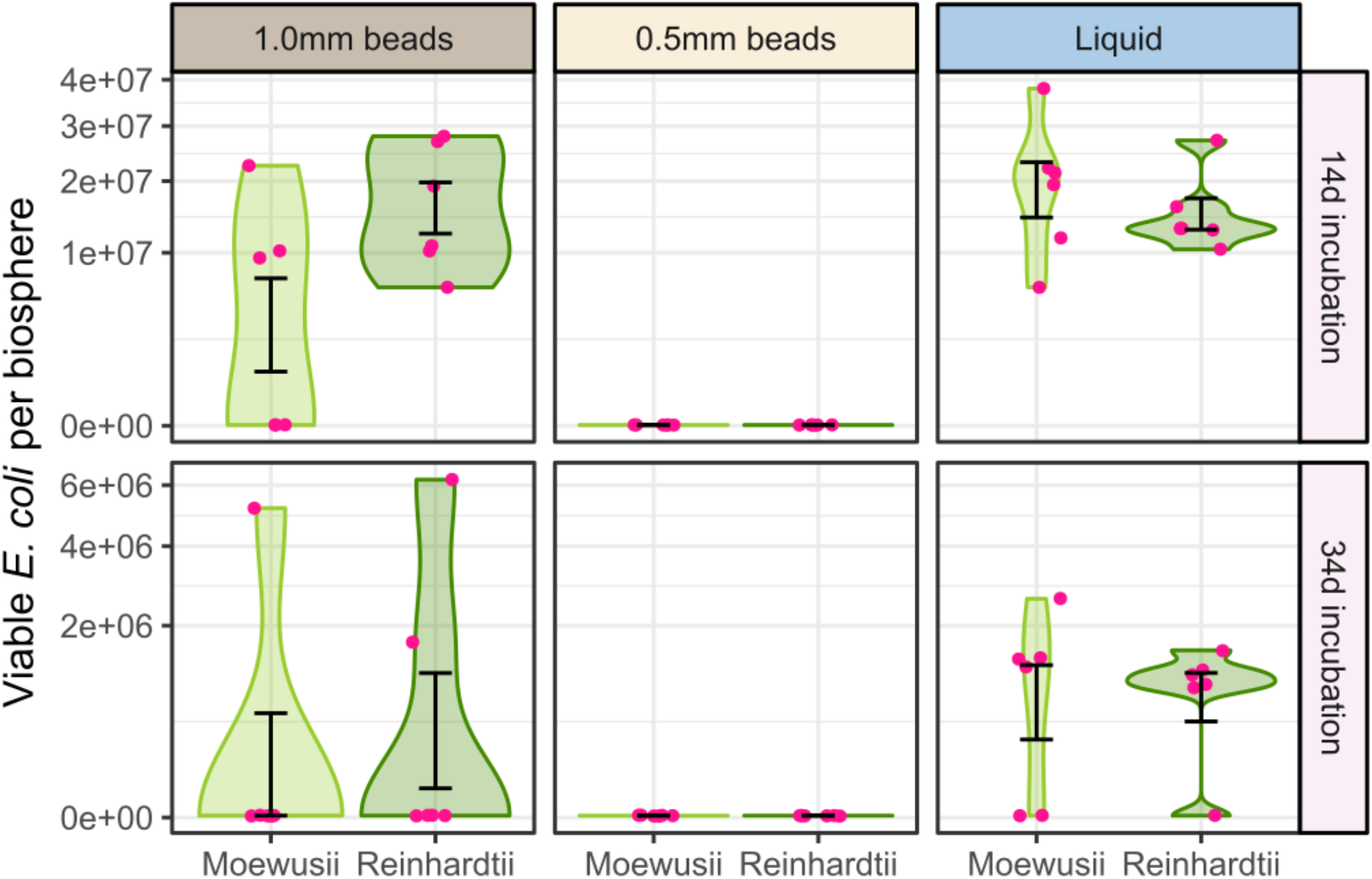
Colony forming units (CFUs) were assessed via plating after 14d (top row) and 34d (bottom row) of incubation. Two microtiter plates each containing three replicates per treatment (n = 6) were destructively harvested at each timepoint. Biospheres were initially inoculated with 2.5 × 10^9^ cells; not shown on plot. Error bars show mean ± standard error. *E. coli* population densities dropped rapidly in biospheres with small-sized substrate, and were reduced, but less dramatically, in biospheres containing larger substrate size and no solid substrate (liquid).

### Experiment 4: Algae cluster around necromass patches

To test our hypothesis that biospheres with smaller substrate particles failed more quickly due to more limited access to resources (necromass), we tracked algal spatial dynamics in biospheres with differing necromass distributions. To do this, we prepared *E. coli* necromass patches equivalent to either 40% (large) or 20% (small) of the target viable biomass that in each biosphere and placed them in the bottom of the biosphere as follows: either one large patch was placed centrally (high heterogeneity treatment), or two small patches were placed on opposing sides of the biosphere well (intermediate heterogeneity treatment). The low heterogeneity resource treatment received a suspension of one large necromass patch resuspended in sterile saline solution, and the ‘no resource’ control received no necromass. All biospheres were filled with 1.0 mm glass beads and inoculated with the CR + EC and CM + EC communities. We then measured chlorophyll autofluorescence daily using a matrix scanning method, allowing higher resolution observation of algal distributions in the biospheres over time.

In this experiment, CR + EC biospheres failed more quickly than in previous experiments (Fig. S5). Despite this, we still observed differences between necromass heterogeneity treatments for both species. Regardless of spatial treatment, all biospheres with the primary producer CR failed within the first week and small but significant differences in survival time were observed: biospheres with the high and low heterogeneity treatment persisted for 7 d (median), which those with intermediate heterogeneity for 6 d, and no resource for 5 d (log-rank test *X*^2^= 25.6, df = 3, p= 1e-05; Fig. S6, Tab. S1). Biospheres with CM as the primary producer, on the other hand, largely persisted during the experimental period (except the ‘no resource’ treatment), and quicker declines in chlorophyll autofluorescence were observed in the intermediate heterogeneity treatment compared to the high and low heterogeneity treatments. Culturable *E. coli* were recovered from relatively few biospheres (55% from CM biospheres and 14% from CR; Tab. S2); however, motile bacteria with the same size and morphology as *E. coli* were observed microscopically. We tested for problems with the media and signs of contamination (Table S3) and further investigated the greater-than-expected loss of culturable *E. coli* replicating these experimental conditions with the CM + EC for a longer experimental duration (31 vs. 13 d). We again recovered culturable cells from fewer biospheres than previous experiments; however, we observed an increasing proportion of biospheres containing culturable cells over time across all treatments (Table S4).

Because of the rapid failure of the CR + EC biospheres, we focused on the CM + EC biospheres for our analysis of algal distribution in the biosphere over time (Fig. 6). We observed that while overall there was a higher chlorophyll autofluorescence reading from the biospheres with a homogeneous necromass distribution, there were higher local concentrations of algae surrounding the necromass patches that developed by day six and were maintained during the 13d experimental period in biospheres containing 1.0mm beads. We tested if fluorescence intensity was stronger in regions of the microtiter well containing a necromass patch by comparing the average fluorescence intensity of the entire well with the average fluorescence intensity within either on central circle of pixels (for the high heterogeneity treatment with one central necromass patch) or semicircles of pixels on the left and right sides of the well (for the intermediate heterogeneity treatment with two necromass patches; Fig. S7).

**Figure 6.**
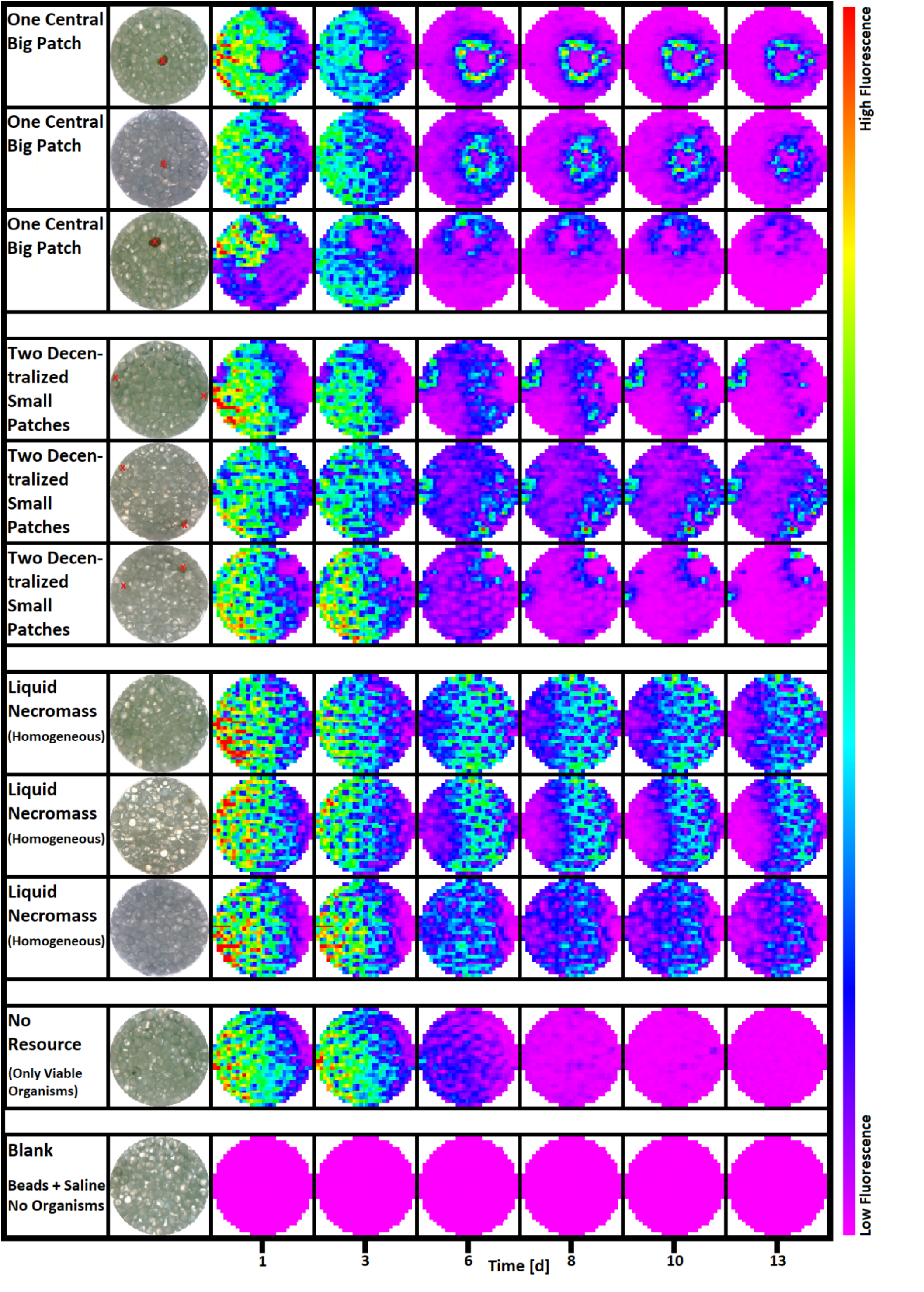
Examples of chlorophyll autofluorescence matrix scans (excitation 470nm, emission 680nm; columns 3-8) from biospheres containing *Chlamydomonas moewusii* and *E. coli* over 13 days. Three examples each are shown for the high heterogeneity treatment (rows 1-3), intermediate heterogeneity treatment (row 4-6) and low heterogeneity treatment (row 7-9). One example each is shown for the no resource control (row 10) and the blank (i.e. containing no living organisms; row 11). Treatment descriptions are provided in column 1 and column 2 shows the spatial location of the necromass patches (in red) when present. *C. moewusii* cells cluster around necromass patches, particularly when one large patch is present (highest resource heterogeneity).

## DISCUSSION

In this work, we show the importance of spatial structure and heterogeneity on population and system persistence in matter-closed microbial biospheres under resource limitation conditions. Substrate particle size dramatically affected survival: biospheres with small substrate particle size failed more quickly than biospheres containing larger substrate particles or no solid substrate particles, and failed to support a decomposer population when the amount of initially inoculated cells to surface area ratio was low (∼30,000 cells mm^-2^; experiments 2 and 3). All effects mentioned above were observed regardless of biosphere liquid volume or primary producer species.

We were surprised at the rapid decline in culturable *E. coli* in the 0.5mm bead treatments (8d in experiment 2 and 14d in experiment 3) because of reports of bacterial species persisting on self-necromass alone for months to years in closed systems (Shoemaker et al., 2021). However, in contrast with their experiment, our systems contained physical substrate particles. Decomposer populations in biospheres with 0.5mm beads likely had limited access to resources to support their growth and reproduction. Spatial structure (manipulated here via plate size, substrate type and size, volume of soil solution) largely determines the distribution of living cells upon inoculation. The location of the living cells at the beginning of the experiment subsequently affects resource distribution: with no other resources available, the microorganisms themselves become the only resource in the system after their death. In biospheres with small (vs. large) substrate particle size, necromass patches would then be distributed in a longer and more tortuous pore network containing more dead ends, necessitating longer travel distances to access comparatively smaller resource pools and more frequent dead ends. These results are in line with a spatially-explicit model that predicts significantly less overall resource consumption in more heterogeneous and less well-connected environments (Nunan et al., 2020). Smaller cell clusters (and therefore smaller necromass patches following cell death) are also predicted in less well-connected environments (Bickel and Or, 2023), which likely strongly limited decomposer growth in the biospheres containing 0.5 mm beads. In addition to limiting microbial motility to necromass resources, soil microstructure is also hypothesized to affect the dispersal of necromass itself (Buckeridge et al., 2022).

In addition to affecting pore network properties, substrate particle size also determines surface area, with biospheres containing 0.5mm beads having approximately twice as much surface area compared to those with 1.0mm beads or sand. Thus our particle size treatments had a large influence on cell density per unit area and distances between cells. The initial cell densities inoculated into our biospheres ranged from 25,000-100,000 cells per square millimeter. These values are within the range observed in temperate forest topsoil aggregates, where the number of neighboring cells (within 20um) reported for cell densities similar to those initially present in our 0.5mm bead treatment were 29, compared to 78 for densities similar to those in our 1.0mm bead treatment (Schmidt et al., 2024). Lower neighbor density translates to less local resources (dead neighbors are a resource), and may have influenced the formation of biofilms, or intraspecific interactions within those that formed (Eigentler et al., 2022).

Spatial distance also affects interspecific interactions: it can decrease competitive pressure (Tecon et al., 2018; Bickel and Or, 2023; Beizman-Magen et al., 2024), and even for species with positive interactions, limited spatial separation promotes higher population densities than co-location (Kim et al., 2008). In producer-decomposer communities, these organisms both compete with and facilitate one another (Daufresne and Loreau, 2001; Loreau, 2001). The loss of decomposer populations in our biospheres led to a system breakdown of this facilitative relationship, evidenced by significantly shorter survival times. While *Chlamydomonas* can acquire nitrogen from some amino acids directly (Calatrava et al., 2019, 2023), it is likely that when both *E. coli* and *Chlamydomonas* are active in our systems that *E. coli* are better competitors for nutrients (Jansson, 1988; Daufresne et al., 2008), and this may in fact be required for system stability (Daufresne and Loreau, 2001). Algae are then largely dependent upon decomposer mineralization of inorganic nutrients, resulting in a lead time until nutrient resources are more available to them: upon initial inoculation no resources are present leading to senescence; nutrients immobilized in this necromass are then largely taken up by decomposers and only after mineralization become available to the algae. In the 0.5 mm bead treatments, larger spatial separation between both the decomposer and necromass, and subsequent mineralized nutrients and the algae may have led to reduced algae and bacteria populations (experiment 1) and the relatively fast collapse of these systems (experiments 2 and 3).

System failure can also be thought about in the context of survivorship (Rillig and Antonovics, 2019), a concept from population ecology that describes the number of surviving individuals over time. In microbial biospherics, the analogue would be the metric of system persistence measured: chlorophyll autofluorescence in our case. Across experiments, systems were characterized by a behavior reminiscent of type III survivorship curves: chlorophyll autofluorescence dropped precipitously during the first days of incubation. Then systems stabilized, declined more gradually before failing, or failed immediately. This type of behavior is hypothesized to occur due to initial system instability (Rillig and Antonovics, 2019). In our experiment, systems sometimes stabilized only when larger substrate particles were present; biospheres containing smaller substrate particles always failed, indicating that they could not overcome the instability imposed by the initially strong resource limitations.

After completing three experiments (1-3) manipulating spatial structure, we hypothesized that access to and the distribution of resources (i.e. realized spatial heterogeneity) could explain the observed differences in system survival. To test this, we created biospheres with explicit differences in necromass (resource) heterogeneity using non-restrictive substrate particle size (i.e. 1.0mm beads) and found that the presence of necromass significantly increased system survival time, and biospheres with either high or low necromass heterogeneity performed better than those with intermediate heterogeneity. Low necromass heterogeneity was generated by adding necromass in suspension, such that the microorganisms were initially co-located with the necromass resource (i.e. everything everywhere) thus allowing for immediate resource access and subsequently higher necromass concentrations throughout the biosphere compared to the other treatments. High heterogeneity on the other hand likely promoted longer persistence times by serving as a central nucleus for metabolic activity. Hotspots of microbial activity surrounding organic material is well documented (Kuzyakov and Blagodatskaya, 2015), and it is likely that both organisms co-occurred around the patch, also dying around the patch, thereby forgoing the high costs of motility (Schavemaker and Lynch, 2022) needed for foraging for resources, and maintaining locally high resource concentrations.

While patterns in performance were similar across all treatments (high and low heterogeneity promoting better system-level performance than intermediate heterogeneity), we observed differences in survival dependent upon the algal partner: biospheres containing *C. reinhardtii* only persisted for 5-7 d while those with *C. moewusii* and necromass persisted for the experimental duration. Therefore, we focused our analysis of algal spatial distribution on the more persistent CM + EC biospheres. We observed a localization of algal populations around necromass patches that developed after 6d of incubation and remained for the duration of the experiment: necromass heterogeneity drove algal spatial organization patterns. Together with the survival results, this provides support for the hypothesis that, in substrates such as soil, spatial heterogeneity alone can explain carbon substrate decomposition (Lehmann et al., 2020).

Together these findings indicate a limit to the spatial/physical parameter space in which producer-decomposer communities can establish and self-sustain via self-recycling of necromass. The findings presented here have implications for microbial necromass recycling in soils, an important process in the pathway of soil carbon storage (Liang et al., 2017) for which the effects of microscale structure are unresolved (Buckeridge et al., 2020). Our results may also help parameterize spatially-explicit models of soil carbon processes. In our experiments, spatial structure was a critical determinant of system failure.

## MATERIALS AND METHODS

Closed biospherics systems were made using 96- or 48-well microtiter plates filled with sterile substrate, inoculated with a mixture of algae (*Chlamydomonas reinhardtii* or *Chlamydomonas moewusii*; producers) and bacteria (*Escherichia coli*; decomposer) suspended in isotonic solution (0.85% NaCl) and sealed with polyolefin sealing foils that were found to minimize water vapor loss, a proxy for gas exchange (de Jesús Astacio et al., 2021). To model a soil-like environment, biospheres were filled to about 1/2 their total volume with substrate and approximately the upper quarter to third of the substrate was not submerged in liquid, depending on particle size. No initial nutrients or media were supplied, so recycling of necromass (the only nutrient resource) was required for biosphere persistence. Biospheres were incubated in a climate-controlled chamber at 25°C with a 16/8h light/dark cycle. To assess system persistence while the biospheres were closed, we measured chlorophyll autofluorescence several times per week. To assess microbial population densities, we performed destructive harvests of some replicate biospheres during the course of each experiment.

### Experiment 1

Four types of differently spatially structured environments were generated using different substrate particle types: 1-2 mm quartz sand, 1 mm glass beads, 0.5 mm glass beads, and no substrate particles (isotonic saline solution only). To test for effects of linear linkages (mimicking the physical spatial connections made by fungal hyphae), ten glass fibers (10 mm, PRECiT) were added to some biospheres containing either sand or 1.0 mm beads. Each biosphere (i.e. single well in a microtiter plate) was inoculated with 50 μL isotonic saline solution containing 1×10^6^ *C. reinhardtii* cells and 1×10^8^ *E. coli* cells (CR + EC community). Using the inner 60 wells three 96-well microtiter plates (Sarstedt) five replicates per treatment per plate (n = 30), plus five replicates each of negative controls (i.e. blanks; no organisms) with sand, 1.0 mm glass beads or saline-only (n = 15) were prepared, for a total of 90 biospheres and 45 blanks. All treatments were present on each plate and were randomly distributed. Biospheres were sealed using ThermaSeal RTS™ (Excel Scientific, Victorville, CA, USA) sealing foils and incubated with multiple chlorophyll auto-fluorescence measurements taken weekly. After 19 days, a destructive harvest was performed and viable *E. coli* and total *C. reinhardtii* cells were assessed by serial dilution spot plating (SD-SPS) and flow cytometry, respectively.

### Experiment 2

To test the effects of substrate size and its interaction with liquid volume on population and system persistence, we modified the design of experiment 1. A fully factorial design was employed: substrate treatments included 1mm glass beads, 0.5mm glass beads and a 50/50 (w/w) mix of 1 mm and 0.5 mm beads (3 levels); volume treatments included the addition of 100 and 200 μL isotonic saline solution (2 levels). Biospheres were inoculated with 2.5×10^6^ *C. reinhardtii* cells and 2.5×10^8^ *E. coli* cells (CR + EC community). Using the inner 24 wells of four 48-well microtiter plates (Greiner Bio-One, Frickenhausen, Germany), three replicates per treatment (n = 18), plus three negative controls (i.e. blanks with no organisms) each containing glass beads and either 100 or 200 uL saline were added to each plate (n = 6), for a total of 72 biospheres and 24 blanks. These were sealed and incubated for 29 days with multiple chlorophyll auto-fluorescence measurements per week. Destructive harvests of two plates (6 replicates per treatment) were conducted after 8 and 29 days of incubation.

### Experiment 3

Loss of viable *E. coli* populations in all 0.5 mm and mixed bead treatments were observed in experiment 2. To confirm these results, and again using the CR + EC community, we replicated two of the spatial structure treatments (1 mm and 0.5 mm beads with 200 μL isotonic saline solution), and additionally applied a no-bead (liquid only) treatment (spatial structure = 3 levels). We also prepared biospheres with all spatial structure treatments using the CM + EC community (community = 2 levels). The same cell densities were used for both communities (2.5×10^6^ algae cells and 2.5×10^8^ *E. coli* cells per biosphere). Using the inner 24 wells of six 48-well microtiter plates (Greiner Bio-One, Frickenhausen, Germany), we set up three replicates per treatment per plate (n = 18), plus three replicates each of negative controls (i.e. blanks; no organisms) with and without 1.0 mm glass beads (n = 6), for a total of 108 biospheres and 36 blanks. Treatments were randomly distributed on each plate. Systems were sealed and incubated for 34 days, with periodic chlorophyll auto-fluorescence measurements. Destructive harvests of two plates (i.e. six replicates per treatment) were conducted after 14 and 34 days of incubation.

### Experiment 4

To explicitly test if the spatial distribution of resource (i.e. necromass) patches could explain the differing population and system persistence observed in experiments 1-3, we generated biospheres differing in the initial spatial location of an *E. coli* necromass patch equivalent to 40% of the biomass of the living biomass (algae + *E. coli*) added to each biosphere. We generated necromass by exposing a pellet of washed *E. coli* (9.17 × 10^8^ cells) to 99.96% ethanol (280 mL) for 3 hours followed by centrifugation, removal of the supernatant and resuspension in 56 mL of 99.96% ethanol; under perfect conditions 200uL of this suspension would contain the target amount of *E. coli* necromass for each biosphere. Assuming a 20% loss of biomass during preparation, either 220 μL (for large necromass patches) or 110 μL (for small necromass patches) were transferred into 0.5 mL microcentrifuge tubes, followed by centrifugation and removal of supernatant. Tubes containing the pellets were left open to dry for 48 hours under sterile conditions. While this necromass production method is quite artificial compared to natural death pathways in soil (Camenzind et al., 2023), it will maintain more biomass building blocks, such as nucleic acids and therefore may be slightly more realistic than autoclaving. After drying, either one large patch or two small patches were added to the bottom of some biospheres. Biospheres were filled with 1mm beads and inoculated with the same communities and used in experiment 3 (CR + EC and CM + EC, community treatment = 2 levels). Roughly double the CR cell density was added compared to CM (Tab. 1), to hold the inoculated biomass constant across biospheres. In addition to the biospheres containing a large necromass patch (high heterogeneity) or two small patches (intermediate heterogeneity), we also added a liquid suspension (one large patch in saline solution) to some biospheres (low heterogeneity; necromass heterogeneity = 3 levels). Using the inner 24 wells of six 48-well microtiter plates (Greiner Bio-One, Frickenhausen, Germany), we set up three replicates per necromass patch X community treatment, and two replicates of a no-necromass control per community (n = 22), plus two blanks (i.e. no organisms), for a total of 132 biospheres and 12 blanks. Treatments were randomly distributed on each plate. Systems were sealed and incubated for 13 days, with periodic chlorophyll auto-fluorescence measurements. Additionally, [explain Clariostar plate scans here] to map the location of algae on the bottom of each plate well and assess their proximity to necromass patches, when present. A destructive harvest of two plates was conducted after 7 days of incubation.

**Table 1.** Overview of experimental conditions.

| Expt | Cell density (cells/well) |  | Biosphere physical conditions |  |  |
| --- | --- | --- | --- | --- | --- |
| | Producer | Decomposer | Substrate(s)<br>(SA cm<sup>2</sup>) | Plate type<br>(tot. vol. μL) | Liq. vol.<br>(μL) |
| 1 | CR: $1 \times 10^6$ | EC: $1 \times 10^8$ | Sand (9.4)<br>1.0mm GB (9.7)<br>0.5mm GB (19.4)<br>None | 96-well<br>(323) | 50 |
| 2 | CR: $2.5 \times 10^6$ | EC: $2.5 \times 10^8$ | 1.0mm GB (51)<br>0.5mm GB (102)<br>GB mix (76.5) | 48-well<br>(1700) | 100 or 200 |
| 3 | CR: $2.5 \times 10^6$<br>or<br>CM: $2.5 \times 10^6$ | EC: $2.5 \times 10^8$ | 1.0mm GB (51)<br>0.5mm GB (102)<br>None | 48-well<br>(1700) | 200 |
| 4 | CR: $4.55 \times 10^6$<br>or<br>CM: $2.5 \times 10^6$ | EC: $2.5 \times 10^8$ | 1.0mm GB (51)<br>± necromass | 48-well<br>(1700) | 200 |

Abbreviations: CR = *C. reinhardtii*; CM = *C. moewusii*; EC = *E. coli*; GB = glass beads; SA = surface area

### Cell culture conditions and cell washing

*Chlamydomonas reinhardtii* (SAG 11-32b) was maintained on yeast-agar slants and in liquid Tris-Acetate-Phosphate (TAP) medium with shaking at 200 rpm and a 16/8 hours light/dark cycle at 25°C in a climate controlled chamber, with weekly transfers into fresh medium. Prior to beginning an experiment, cells were transferred into fresh media to a density of 10^5^ cells per mL and incubated for five days. Live cell density was determined microscopically at 200X with a hemocytometer and Evan’s blue dye. *E. coli* MG1655 (DSM 18039) stock cultures were maintained in Luria Broth (LB) medium with 25% glycerol and stored at -70°C. Prior to the beginning of an experiment, a small amount of stock culture was transferred into fresh LB medium and incubated for 18h at 37°C at 180 rpm. Cell density was determined by measuring the blank corrected absorbance at 600 nm (OD600) and compared to a standard curve. Known quantities of cells were washed of media before inoculation into biospheres by centrifugation at 3000 x g for 5 minutes, removal of supernatant and resuspension in 0.85% saline solution. Cell washing was repeated twice before resuspending the pellet to the target density in saline.

### Tracking data collection: chlorophyll auto-fluorescence measurement in sealed biospheres

We assessed algal viability in the closed systems via chlorophyll auto-fluorescence intensity measurements using a microplate reader (FLUOstar Omega, excitation 470nm, emission 680nm). The reported value per well is the average of twenty fluorescence measurements in a circular pattern; the same gain setting was used across all measurements within one experiment. Because the plates contained opaque or semi-transparent substrate particles we could not measure the entire biosphere. To measure a representative proportion of the algae population, microtiter plates were covered and incubated over light for 2 hours before measurements to encourage phototropic movement of algae to the bottom of the plate wells, where the measurement was taken. Loss of the chlorophyll auto-fluorescence signal (i.e. measurements below the detection limit) was used as an indication of system failure: without photosynthetic activity, all respired CO_2_ would be lost to the atmosphere with no re-uptake mechanism and therefore could not self-sustain (Rillig and Antonovics, 2019). In previous experiments we observed a rapid decline in fluorescence intensity during the first 7-10 days of the experiment followed by either a stabilization or loss of the signal. Therefore we conducted measurements more frequently during the first week of the experiments (5-7 times per week) and less frequently afterwards (2-3 times per week).

### Endpoint data collection: destructive cell recovery and cell enumeration

While the non-invasive tracking measurements provide data about the algal viability in the systems, destructive harvests are necessary to gain information about the bacterial and algal population sizes. Once a biosphere is opened, the experiment is considered concluded because of the immediate mixing of the atmosphere in the room with the biosphere atmosphere. In experiment 1 a destructive harvest was conducted after 19 d incubation, for experiments 2-4, destructive harvests were conducted once or twice per experiment: once shortly following loss of fluorescence signal by microplate reader, and sometimes again after several more weeks of incubation. Under sterile conditions, plates were tapped against the bench to collect condensation gathered on the sealing foil and opened. Isotonic saline (0.85% NaCl) solution (200 μL) was added to each biosphere, mixed thoroughly by pipetting up and down, and 200 μL were recovered from each biosphere for enumeration. A 10 μL aliquot of this was then used to prepare a serial dilution which was then plated on LB-agar plates to enumerate viable *E. coli* cells, using the single plate-serial dilution spotting (SP-SDS) method (Thomas et al., 2015) to plate seven dilution levels on each plate. An additional aliquot of the biosphere recovery solution was mixed with additional recovery solution to prepare a dilution for flow cytometry (Guava^®^ easyCyte™, Cytek Biosciences, Fremont, CA, USA) to enumerate total and fluorescing algal cells. Settings for each species were standardized using a positive control sample to exclude e.g. cell debris and 5000 events were collected per sample.

### Statistics

Data analysis was performed using R version 4.4.0 (R Core Team, 2021). Chlorophyll autofluorescence (i.e. tracking) data was used to assess the persistence of closed systems and determine time to system failure. Resulting from background fluorescence of many materials, the fluorescence intensity value in the absence of chlorophyll is not equal to zero. Therefore we used the background fluorescence readings of ‘blank’ wells (containing sterile substrate particles and isotonic solution, but no organisms) to calculate a limit of detection threshold equivalent to the mean plus three standard deviations of the blank values. For each experiment, blanks were prepared with the same substrate particles as the experimental treatments. Once the chlorophyll autofluorescence readings from a biosphere fell below this threshold, it was considered to have failed. Using these data, we determined the first date of failure (i.e. chlorophyll auto-fluorescence value below the detection limit) for all biospheres for which this event occurred. Because not all biospheres failed during the duration of each experiment, biosphere persistence time data was right censored. Therefore we employed survival analysis to compare biosphere persistence times between treatments using the “survival” package (Clark et al., 2003; Therneau, 2024), including median survival time and testing for differences in survival between groups using the log-rank test. Kaplan-Meier plots were used to visualize survival probability for each treatment group via the “ggsurvfit” package (Sjoberg et al., 2023). Differences in population densities (cells per biosphere) of cells recovered from biospheres at harvest were tested using the Kruskall-Wallis test, and when differences were observed, Dunn’s post hoc test. Data visualizations were produced using the R package “ggplot2” (Wickham, 2016).

## ACKNOWLEDGEMENTS

We thank Vera Engelhardt, Tim Bliß and Kari Oelke for support with data collection and methods optimization, and Florian Heyd and Felix Oswaldt for support with the flow cytometry measurements.

