## Supplementary Material for "Spatial structure affects the establishment and persistence of closed microbial ecosystems"

**S1. Experiment 1 supplemental data**

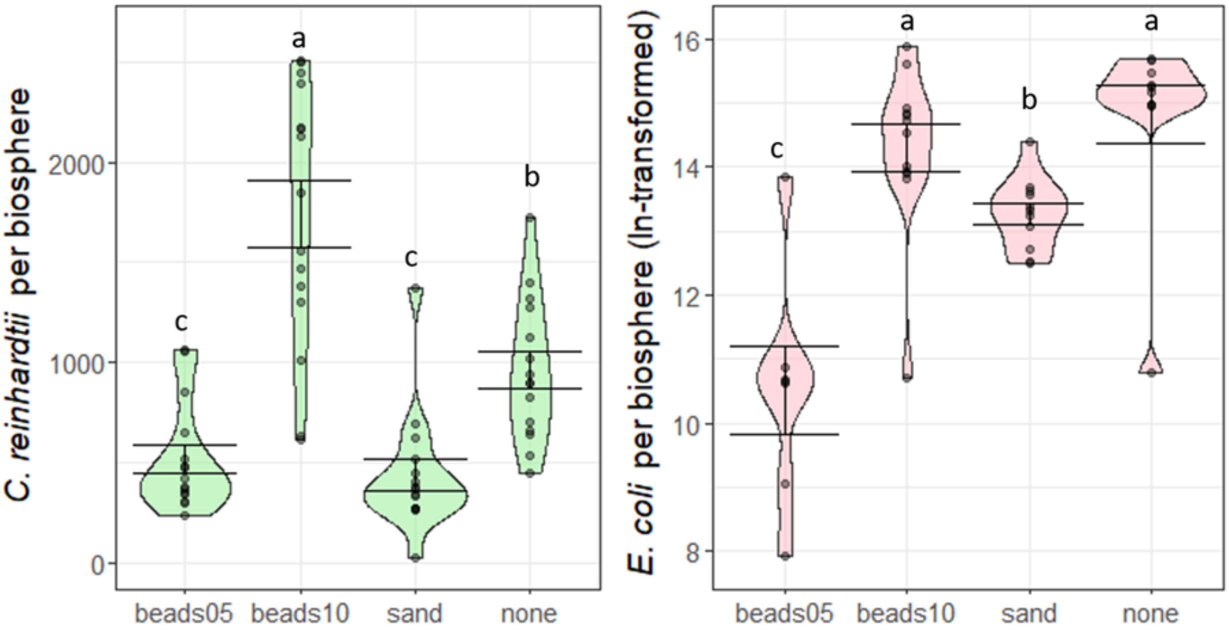


Figure S1. Substrate size and type affects microbial population size. Population densities per biosphere of total *C. reinhardtii* cells (assessed by flow cytometry, L) and viable *E. coli* cells (R, ln-transformed) after 19d of incubation in biosphere systems with different substrate types: beads05 = 0.5 mm beads, beads10 = 1.0 mm beads, sand = 1-2mm quartz sand, none = no substrate addition. Error bars indicate mean +/- standard error. Different letters above violin plots indicate groups that significantly differ (Dunn test with Bonferroni correction, $p < 0.05$).

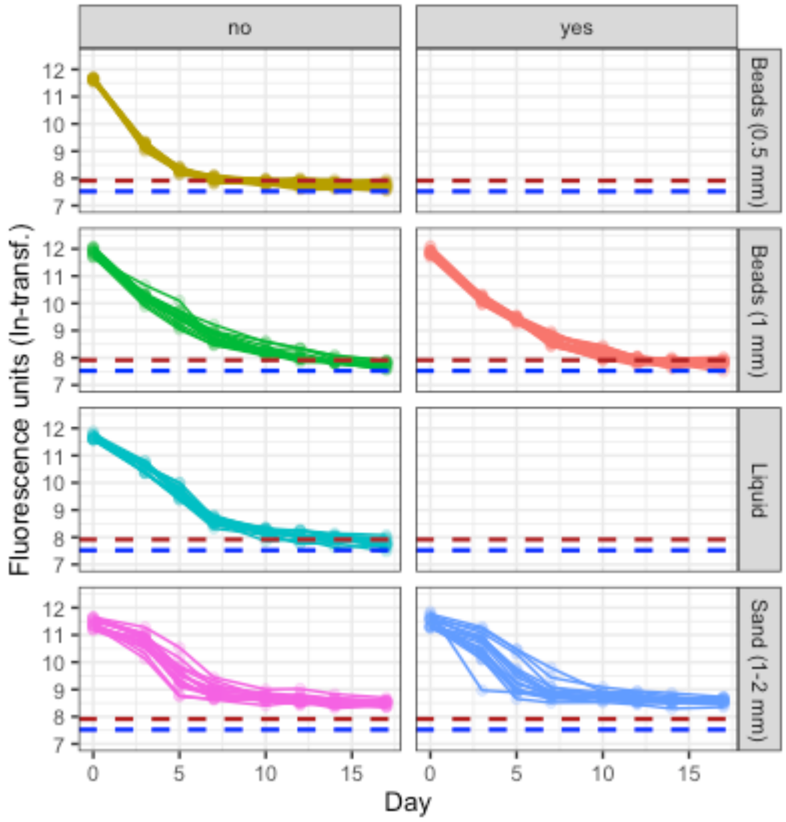


Figure S2. Biosphere persistence time varies by spatial structure/substrate type, but is not affected by the presence of spatial linkages (glass fibers; treatments with glass fibers added are indicated by “yes” in the right-hand column). Each line represents the natural log-transformed chlorophyll autofluorescence measured for a single biosphere over the course of the experiment. The red and blue dashed lines indicate the thresholds for system failure (i.e. the detection limit) for units containing sand or glass beads (red) or liquid-only (blue). Background fluorescence was lower in biospheres with no substrate (liquid only) therefore that treatment has a lower threshold value.

Glass fibers were added to some biospheres (along with either sand or 1.0mm glass beads) to mimic the spatial linkages in soil created by the presence of fungal hyphae. No effects of the glass fiber treatment were observed, therefore data from these biospheres has been excluded from the main text.

**S2. Experiment 2 supplemental data**

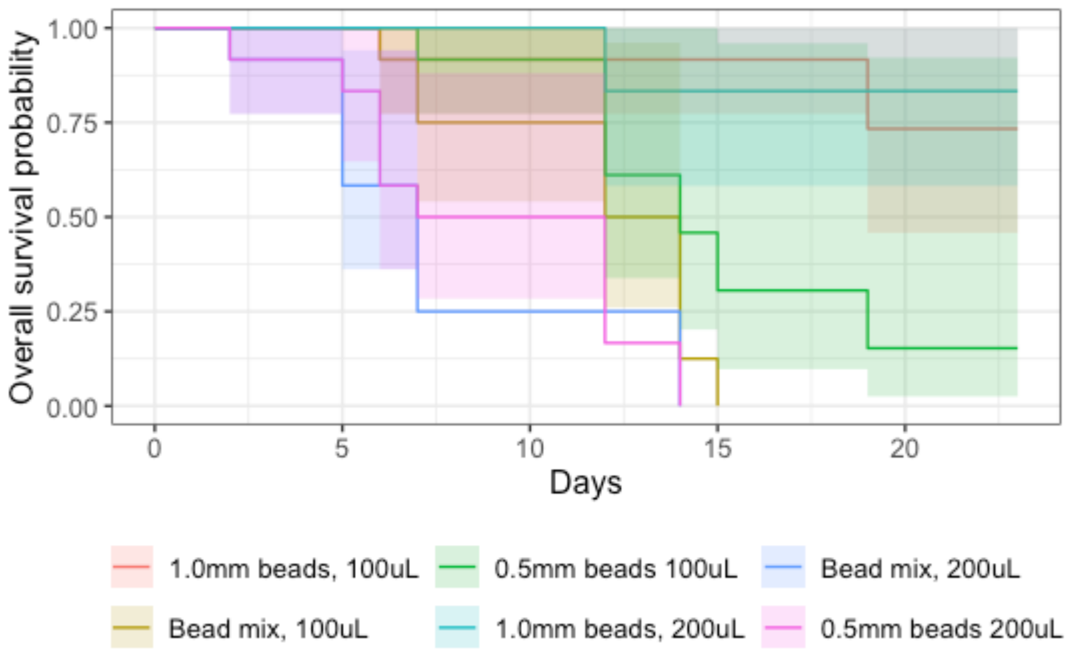


Figure S3. Biospheres with small substrate (0.5mm beads) have lower overall survival probability compared to those containing larger substrate (1.0mm beads). Kaplan-Meier plot showing differences in overall survival probability over time between biospheres containing different substrates (1.0mm beads, 0.5mm beads and a 50/50 mix), and different volumes (100 or 200μL).

### S3. Experiment 4 supplemental data

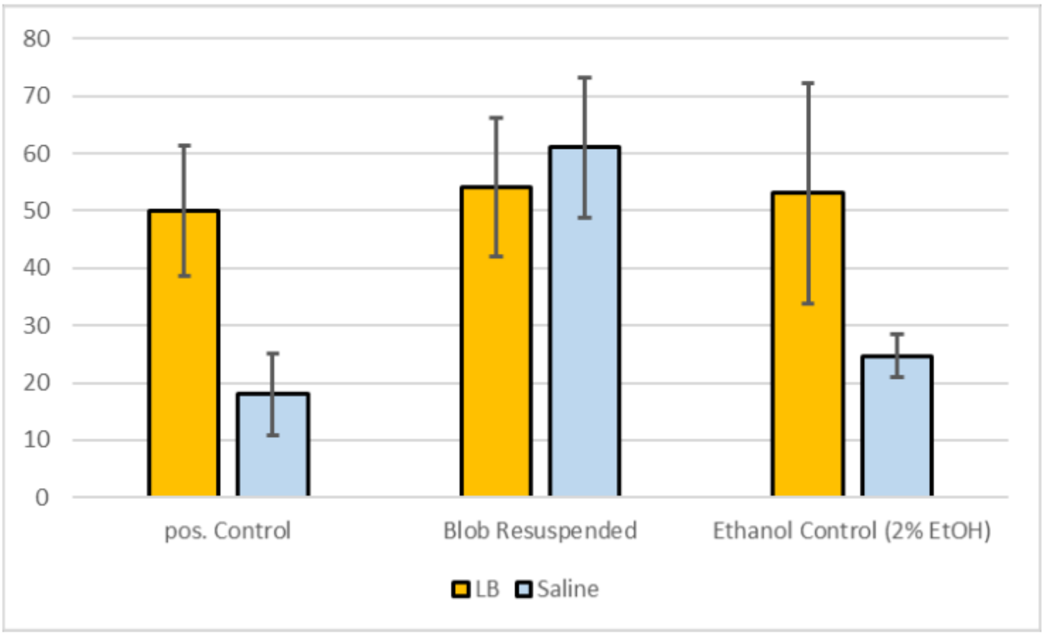


Figure S4. Necromass patches did not negatively impact *E. coli* growth after 60h in either Luria Broth (LB) medium or isotonic saline solution (0.85% NaCl). “pos. Control” = no addition of necromass patch, “Blob Resuspended” = addition of necromass patch, “Ethanol Control” = addition of the 99.8% ethanol (final concentration = 2%) used to produce the necromass patches.

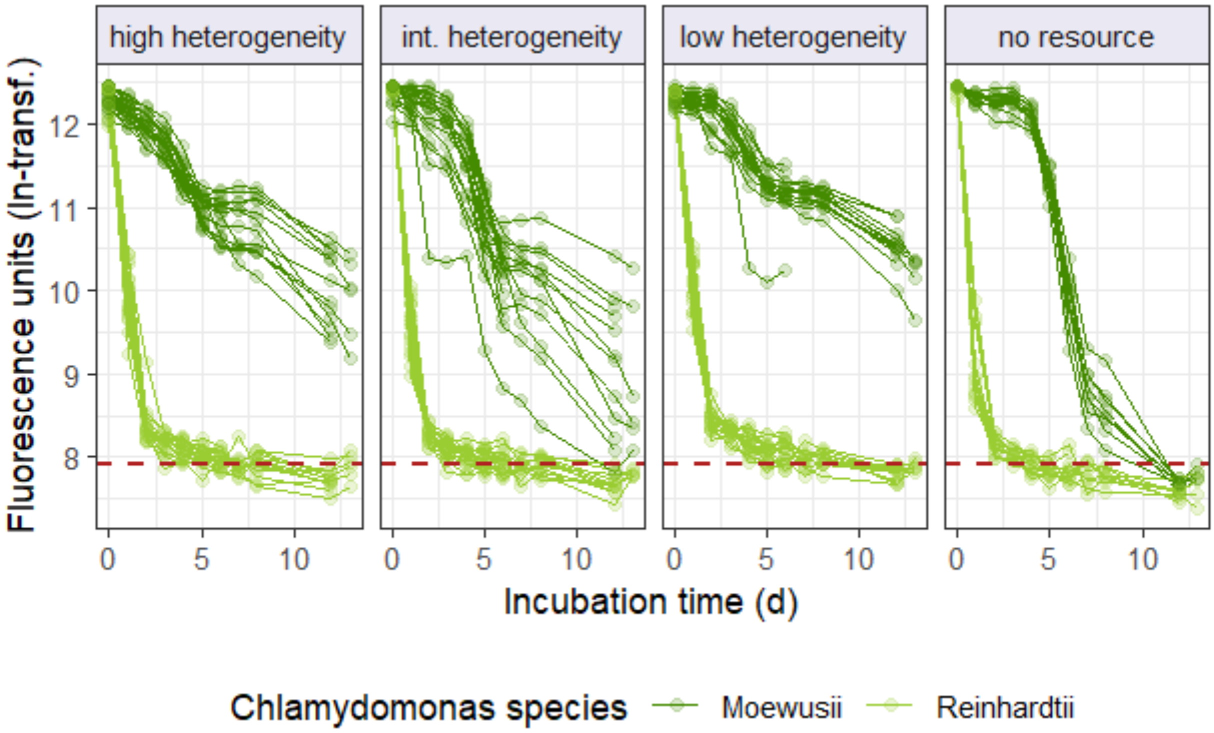


Fig. S5. Biospheres with either high or low necromass heterogeneity persist longer than those with intermediate heterogeneity. Each line represents the natural log-transformed chlorophyll autofluorescence measured for a single biosphere over the course of the experiment. The red dashed line indicates the threshold for system failure (i.e. the detection limit). Biospheres containing *C. moewusii* as the primary producer are shown in dark green, those with *C. reinhardtii* in light green.

Biospheres containing *C. reinhardtii* failed more quickly than those containing *C. moewusii*. Among the *C. moewusii* biosphere, those with no added resource failed within the 13d duration of the experiment. Chlorophyll autofluorescence in the intermediate heterogeneity treatment declined more quickly than in the low or high heterogeneity treatment.

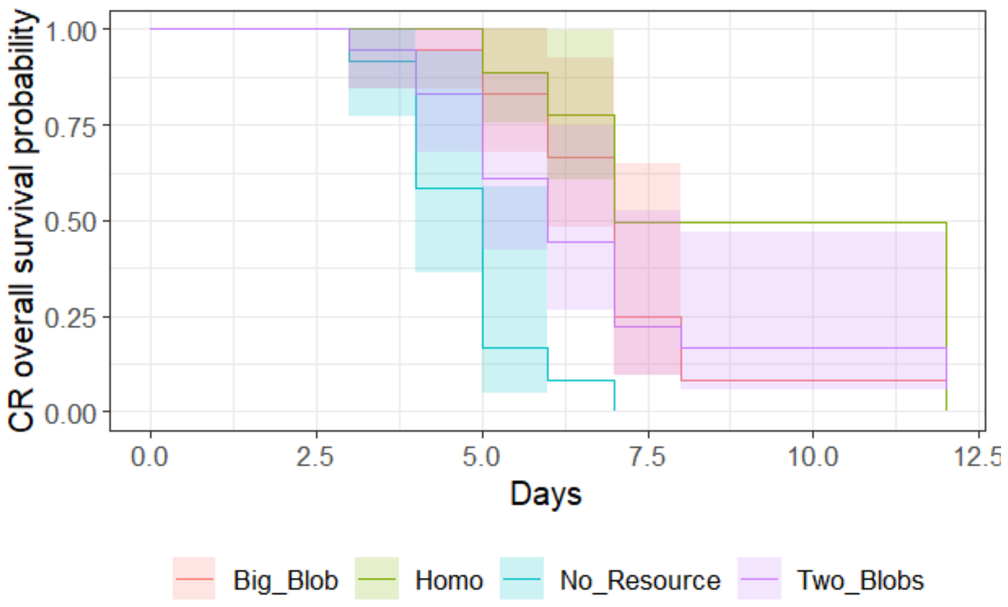


Figure S6. Kaplan-Meier plot showing differences in persistence times between necromass heterogeneity treatments in biospheres with *C. reinhardtii* as the algal partner.

Table S1. Pairwise comparisons of survival in biospheres containing *C. reinhardtii* using Log-Rank test, p value adjustment method: Benjamini-Hochberg

| | High | Low | No resource |
|---|---|---|---|
| Low | 0.06675 | - | - |
| No resource | 0.00051 | 3.5e-05 | - |
| Intermediate | 0.60066 | 0.06662 | 0.01981 |

Table S2. Experiment 4: percentage of biospheres containing culturable *E. coli* after 7d incubation.

| | Algal partner | |
|---|---|---|
| Heterogeneity treatment | *C. moewusii* | *C. reinhardtii* |
| High | 50% | 0% |
| Intermediate | 50% | 16.7% |
| Low | 83% | 33.3% |
| No necromass control | 25% | 0% |
| Total | 54.5% | 13.6% |

Table S3. Tests conducted to investigate discrepancy in *E. coli* results between experiments 2/3 and 4

| Test | Question/method | Result |
|---|---|---|
| Method optimization: spot plating | Will using a larger volume of solution recovered from the biosphere improve detection? | no |
| Media | Was the LB media used defective or otherwise inhibiting growth? | no |
| | Do other colonies appear on R2A agar? | no |
| Microscopy | Do we observe any bacterial cells with a different size and/or morphology than *E. coli*? | no |

Table S4. Percentage of biospheres containing culturable *E. coli* in repetition of experiment 4 (CM + EC community only)

| | Incubation time | | |
|---|---|---|---|
| Heterogeneity treatment | 8d | 16d | 31d |
| High | 20% | 70% | 90% |
| Intermediate | 10% | 50% | 50% |
| Low | 40% | 80% | 77.8% |
| No necromass control | 0% | 66.7% | 83.3% |
| Total | 19.4% | 66.7% | 72.2% |

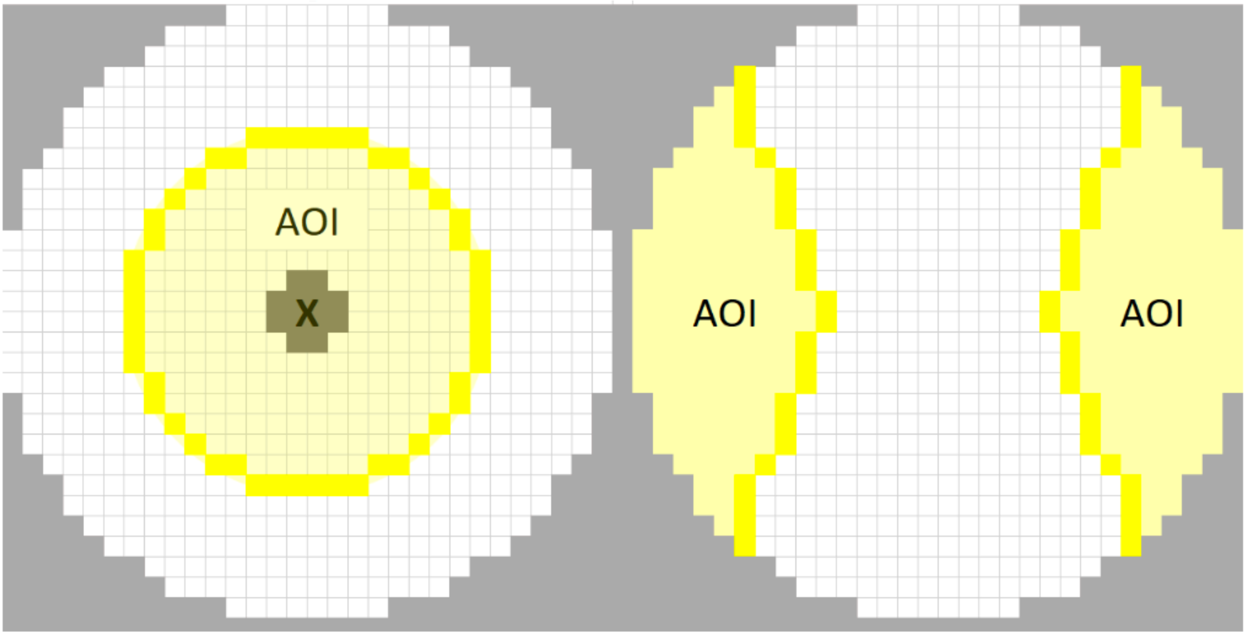


Figure S7. Pixels used in comparison tests to determine if higher than expected fluorescence intensity occurred in regions with a necromass patch for the high heterogeneity treatment (L) and intermediate heterogeneity treatment (R).
